# HDAC inhibitor treatment restores transcriptome, chromatin accessibility and memory deficits in a mouse model for Down syndrome

**DOI:** 10.64898/2026.03.20.712945

**Authors:** Cesar Sierra, René A. J. Crans, Mara Dierssen

## Abstract

Down syndrome (DS) is associated with widespread molecular alterations, including aberrant gene expression, which not only affects chromosome 21 genes but perturbs the whole transcriptome. The genome-wide nature of this alteration suggests that it might be epigenetically mediated, which may also offer therapeutic opportunities given the reversibility of epigenetic modifications. Here, we show that decreased global acetylation level in the hippocampus of the trisomic mouse model Ts65Dn is associated with decreased chromatin accessibility at gene promoters in hippocampal neurons. Strikingly, pharmacological restoration of histone acetylation using the clinically approved histone deacetylase inhibitor suberoylanilide hydroxamic acid (SAHA) normalized transcriptomic, epigenetic and cortico-hippocampal memory deficits in trisomic mice. This rescue was mediated by an unexpected heterochromatization of the supernumerary chromosome. Altogether, our results identify an unanticipated epigenetic mechanism linking histone acetylation dynamics to chromosomal dosage homeostasis.

## Introduction

Down syndrome (DS or trisomy 21) results from an extra copy of human chromosome 21 (HSA21) and is the leading genetic cause of intellectual disability, affecting over 5 million individuals worldwide. The condition profoundly affects brain development and function, leading to impaired cognition and adaptive behavior. Among the most characteristic neurological features of DS, deficits in hippocampal-dependent learning and memory^1^, accompanied by molecular and cellular abnormalities have been identified both in individuals with DS and in mouse models^2–4^. Beyond gene-dosage effects of the triplicated chromosome, DS is characterized by genome-wide alterations in gene expression that contribute to the complex etiology of this disorder^4–7^. Several HSA21 genes, such as *DYRK1A*, *ETS2*, *HMGN1*, *DNMT3L* and *RUNX1* are involved in epigenetic regulation, thereby affecting transcription across the genome. Consistent with this view, multiple studies have reported evidence of chromatin dysfunction in DS, supporting the idea that trisomy perturbs higher-order chromatin organization and transcriptional regulation at a genome-wide scale^8–13^.

Among epigenetic modifications, histone acetylation has been strongly associated with memory formation and consolidation. Acetylation of histone H3 promotes chromatin accessibility and the transcription of genes required for long-term memory storage^14,15^. Supporting this role, pharmacological studies have shown that histone deacetylase inhibitors (HDACis) facilitate memory formation^16–19^, whereas inhibition of histone acetyltransferases (HATs) impairs it^20–23^. Furthermore, histone acetylation alterations have been described in several models of neurological disorders with cognitive dysfunction^24–27^, such as the Ts65Dn mouse model of DS, in which we previously showed reduced hippocampal acetylation of histone H3 (H3K9/K14)^11^. Such hypoacetylation may contribute to the transcriptional dysregulation observed in DS and may interfere with the induction of gene expression programs required for memory formation and maintenance^28^. Given the dynamic and reversible nature of histone acetylation, controlled by the activity of histone acetyltransferases and deacetylases, pharmacological manipulation of this pathway represents an attractive therapeutic strategy for cognitive disorders.

Histone deacetylase inhibitors, including the clinically approved compound suberoylanilide hydroxamic acid (SAHA, also known as vorinostat), increase histone acetylation and modulate chromatin accessibility and transcriptional activity. Based on the view that histone hypoacetylation represses the transcription of plasticity-related genes, HDAC inhibition would be expected to release transcriptional repression and enhance the expression of memory-related genes. However, the impact of HDAC inhibition on the complex transcriptional imbalance associated with chromosomal aneuploidy remains poorly understood. In particular, it is unclear whether epigenetic therapies designed to enhance transcription might exacerbate the gene dosage imbalance inherent to trisomy.

In the present study, we demonstrate in a neuron-specific manner that hippocampal neurons in Ts65Dn exhibit genome-wide epigenetic and transcriptional alterations, associated with global histone hypoacetylation and reduced chromatin accessibility at gene promoters. Surprisingly, pharmacological restoration of histone acetylation using SAHA did not simply increase gene expression programs associated with memory. Instead, SAHA treatment normalized the expression of trisomic genes across the supernumerary chromosome, revealing an unexpected regulatory effect on gene dosage imbalance. This was accompanied by a complete rescue of object recognition memory. Our analyses indicate that HDAC inhibition induces heterochromatinization of the supernumerary chromosome, presenting an atypical underlying molecular mechanism. Together, these findings uncover a previously unrecognized epigenetic mechanism linking histone acetylation dynamics to the regulation of chromosomal dosage effects.

## Results

### Trisomy leads to genome-wide changes in chromatin accessibility and transcription in hippocampal neurons

We characterized the transcriptional and chromatin accessibility changes associated with the trisomy (TS) using assay for transposase-accessible chromatin (ATAC-seq) and RNA-seq analyses in hippocampal neuronal nuclei isolated from trisomic and euploid littermates by Fluorescent-Activated Nuclear Sorting (FANS) (Fig. 1A and Suppl Fig 1A).

**Figure 1.**
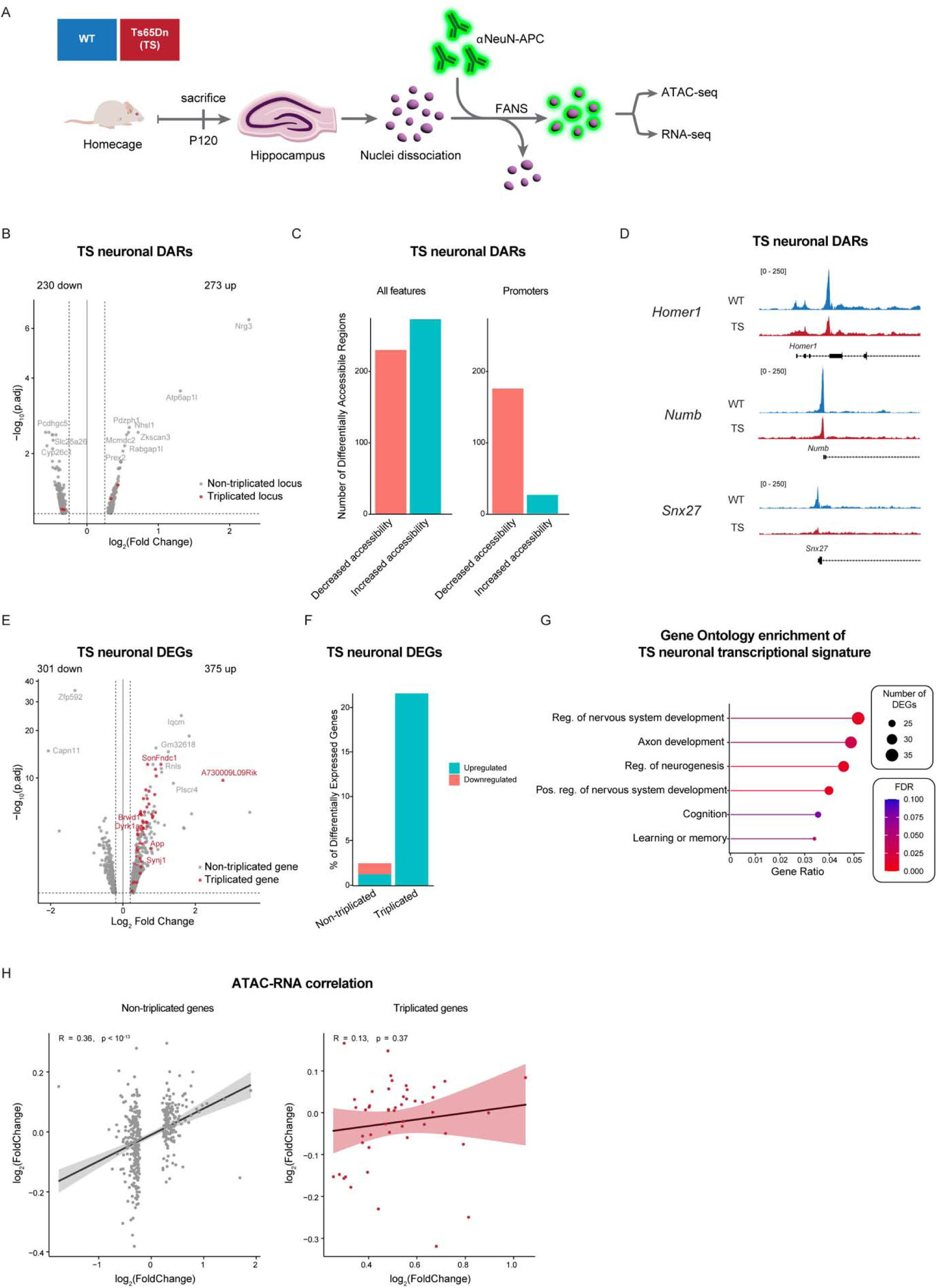
Genome-wide accessibility and transcriptional changes. **A.** Outline of the experimental strategy. **B**. Volcanoplot of detected DARs. DARs located in the triplicated chromosome are colored in red. The name of the closest gene to the peak is depicted, as well as the total number of regions with gained and lost accessibility. Dashed lines indicate the significance thresholds for log2FC and p.adjusted values. **C**. DARs separated by increasing vs. decreasing accessibility regions for all features (left) and for promoters-only (right). **D.** ATAC signal IGV tracks of examples of differentially accessible promoters on genes related to neuronal function. **E.** Volcanoplot of detected DEGs. DEGs located in the triplicated chromosome are colored in red. Number of up- and downregulated genes is indicated. Dashed lines indicate the significance thresholds for log2FC and p.adjusted values. **F.** Barplot showing the proportion (in %) of differentially expressed genes by their triplication status. **G.** Top biological pathways enriched for DEGs identified in TS hippocampal neurons. **H.** Correlation plots between ATAC and RNA signals for non-triplicated genes (left) and triplicated genes (right).

For ATAC-seq, after identifying high-confidence open chromatin regions, the principal component analysis (PCA) showed that samples were separated by genotype in PC1, suggesting a trisomy signature on the epigenetic landscape (Suppl Fig. 1B). Consistently, we identified a total of 503 differentially accessible regions (DARs; FDR<0.1, log2(FoldChange)>0.25; Fig. 1B, Table S1) distributed across multiple chromosomes (Suppl Fig 1C). Genomic annotation of these peaks showed that gene promoters, defined as regions within 3 kb of a transcription start site (TSS), constituted a large portion of the DARs (Suppl Fig. 1D). While the total number of regions with increased and decreased accessibility was comparable in TS neurons, promoter-associated DARs showed a pronounced bias toward reduced accessibility in TS neurons relative to euploid controls (Fig. 1C). This finding is consistent with our previously reported decreased H3K9/K14 acetylation, which is enriched at open gene promoters. Notably, TS neurons showed an increased chromatin compaction on the promoter region of *Homer1*, which encodes for a family of postsynaptic density proteins with key roles in the control of synaptic plasticity^29,30^ (Fig. 1D). Similarly, *Snx27* and *Numb*, which encode for proteins involved in synaptic plasticity and learning^31,32^ also showed a decreased chromatin accessibility at their promoters.

To investigate how this genome-wide epigenetic alteration impacts the transcriptome, we performed RNA sequencing (RNA-seq) on the same sorted neuronal population. PCA analysis revealed a clear separation of samples by genotype (Suppl. Fig. 1E), indicating that trisomy broadly reshapes the neuronal transcriptome. Differential expression analysis identified 676 differentially expressed genes (DEGs, FDR<0.05; Fig. 1E, Table S2). As observed for DARs, transcriptional changes were distributed across the whole genome (Suppl Fig. 1F). Nevertheless, upregulated genes showed a clear enrichment in the triplicated chromosome (Fig 1F, Suppl. Fig. 1F and 1G). Gene Ontology (GO) analysis of DEGs revealed a marked enrichment for categories related to neuronal function, including cognition, learning, and memory (Fig. 1G, Table S3).

Lastly, integration of ATAC-seq and RNA-seq datasets demonstrated a significantly positive correlation between changes in chromatin accessibility and gene expression (Fig. 1H, left), supporting a causal role for chromatin remodeling in driving genome-wide transcriptional alterations in trisomic neurons. In contrast, the absence of a correlation between chromatin accessibility and expression levels of triplicated DEGs (Fig. 1H, right) suggests that their upregulation is primarily driven by gene dosage effects resulting from the additional chromosomal copy.

### HDACi rescues recognition memory impairment in TS mice

Given the reversible nature of epigenetic marks, including histone acetylation, we tested whether increasing histone acetylation through the chronic administration of the histone deacetylase inhibitor (HDACi) suberoylanilide hydroxamic acid (SAHA) could recover the expression of memory-related genes. TS mice received SAHA dissolved in drinking water for four weeks, and an independent group of TS animals was administered a vehicle (HP-β-CD in drinking water) (Fig. 2A).

**Figure 2.**
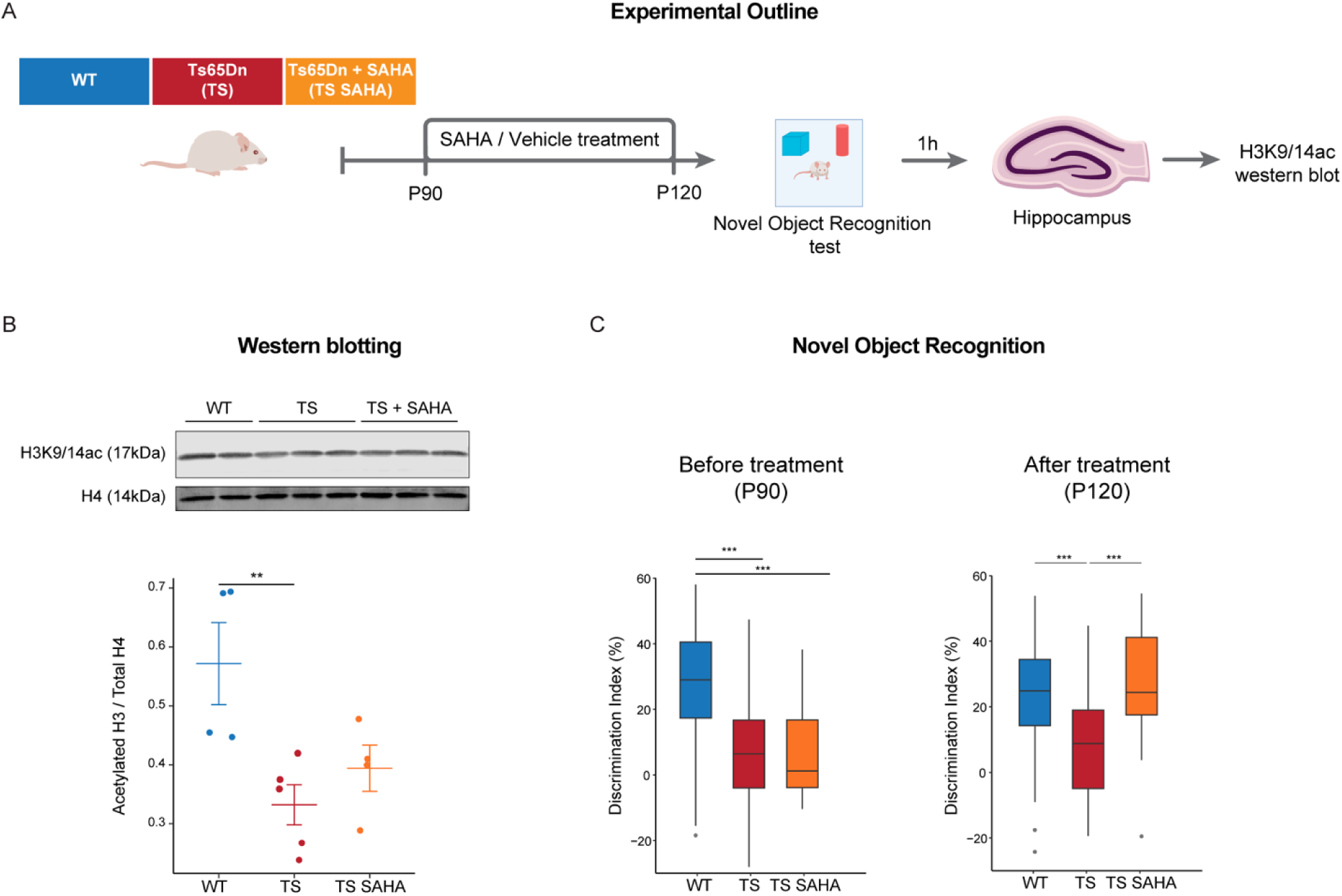
SAHA treatment partially rescues histone hypoacetylation and restores memory performance in NOR. **A.** Outline of the experimental design. **B**. Purified histone proteins from hippocampi immunoblotted for AcH3K9/K14 and total H4 in WT and TS with and without treatment with SAHA (top) and densitometric ratios of AcH3K9/K14 normalized to total H4 (bottom). N = 4/5 per group. Kruskal–Wallis test followed by pairwise Wilcoxon tests with multiple-comparison correction * p < 0.05. **C.** NOR performance before (left) and after (right) SAHA treatment. WT N = 40, TS N = 30, TS + SAHA N = 30. Two-way ANOVA, Tukey HSD as Post-hoc. *** p < 0.001.

Vehicle-treated TS mice exhibited a significant reduction in hippocampal H3K9/K14 acetylation levels compared to WT controls (Fig. 2B). In contrast, SAHA-treated TS animals showed a partial but significant restoration of H3K9/K14 acetylation.

To determine whether this epigenetic rescue translated into functional improvements in cognition, we assessed hippocampus-dependent recognition memory using the Novel Object Recognition (NOR) task. Owing to its non-aversive nature and lack of carryover effects, NOR was performed both before and after treatment. At baseline, WT mice robustly discriminated against the novel object, whereas TS mice failed to exhibit object recognition (Fig. 2C), as previously reported^33,34^. Strikingly, following SAHA treatment, trisomic mice displayed a complete rescue of recognition memory, performing at the same level of WT animals.

Importantly, the impact of the SAHA treatment was specific to the cognitive function, and did not induce changes in other behavioral variables including thigmotaxis, locomotor activity and explorative behavior (Suppl Fig. 2A).

### HDAC inhibition remodels chromatin accessibility and restores global transcriptional programs

After the observation that the rescue of histone hypoacetylation mediated by SAHA was only partial, we aimed at uncovering the molecular changes underlying the rescuing effects of SAHA on recognition memory. Importantly, in this experiment all animals were sacrificed 1 h after completing the NOR task, allowing us to assess the combined effects of genotype and SAHA treatment in the post-learning state (Fig. 3A). This design differs from the experiment shown in Figure 1, where chromatin accessibility and transcription were analyzed in animals maintained in the home-cage condition without behavioral stimulation, providing a baseline genotype signature. We sorted the neuronal nuclei from WT, untreated TS (TS) and SAHA-treated TS (TS-SAHA) mice by FANS 1h after completing the NOR test and performed ATAC-seq and RNA-seq in parallel (Fig. 3A). Thus, all samples analyzed in this experiment correspond to post-learning animals, allowing us to assess the combined effects of genotype and SAHA treatment after the behavioral task.

**Figure 3.**
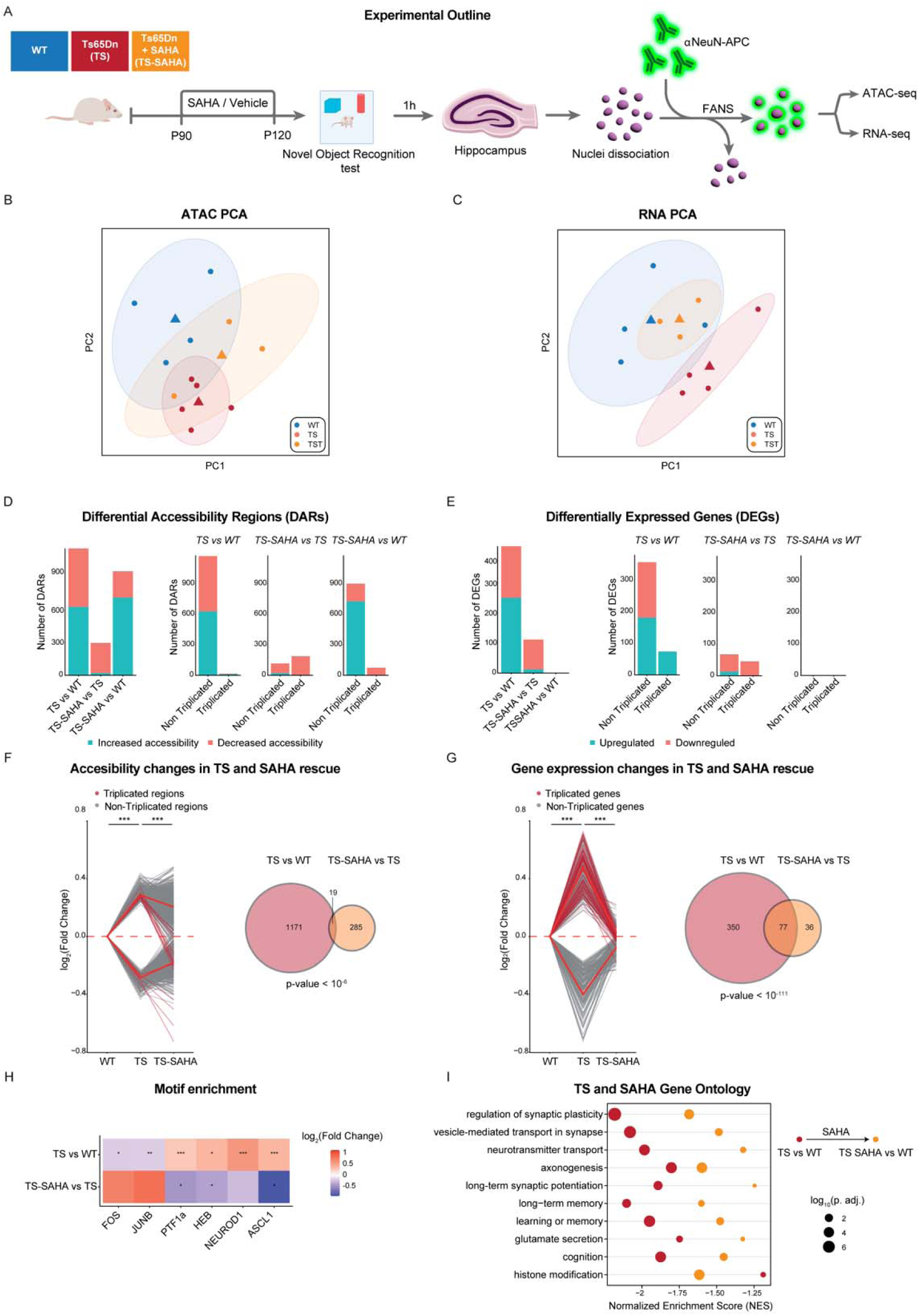
SAHA rescues chromatin and transcriptomic changes. **A.** Outline of the experimental strategy. **B**. PCA plot of ATAC-seq data obtained from four WT, five TS mice and three TS-SAHA (TST). **C.** PCA plot of RNA-seq data obtained from four WT and four TS mice and three TS-SAHA (TST). **D.** Number of differentially accessible regions (DARs) for each comparison (left), and number of DARs in non-triplicated and triplicated genomic regions (right). **E.** Number of differentially expressed genes (DEGs) for each comparison (left) and number of DEGs in non-triplicated and triplicated genomic regions (right). **F.** Analysis of differentially accessible regions (DARs) in TS neurons and their rescue upon SAHA treatment, shown as line-plots (left) and Venn diagram (left). Kruskal–Wallis test followed by pairwise Wilcoxon tests with multiple-comparison correction *** p < 0.001 **G.** Analysis of differentially expressed genes (DEGs) in TS neurons and their rescue upon SAHA treatment, shown as line-plots (left) and Venn diagram (right). Kruskal–Wallis test followed by pairwise Wilcoxon tests with multiple-comparison correction *** p < 0.001 **H.** Heatmap of transcription factor motif enrichment among gained peaks in TS and TS-SAHA neurons, with significance and fold enrichment. **I.** Gene set enrichment analysis of differentially expressed genes in TS (red) and TS-SAHA neurons (orange).

PCAs revealed a clear separation between untreated WT and TS that underwent the learning task in both ATAC-seq and RNA-seq datasets (Fig 3B-C). Notably, TS-SAHA partially clustered with WT in the ATAC-seq PCA and almost completely overlapped with WT in the RNA-seq PCA. In line with these observations, many DARs (Fig. 3D) but no DEGs (Fig. 3E) were identified when comparing TS-SAHA with WT animals.

At the chromatin level, the genome-wide accessibility changes (Fig. 3D, Suppl Fig. 3A-B, Table S4) were similar to those detected in TS animals that did not undergo a learning task (Fig. 1B) indicating that the genotype-driven chromatin signature is largely preserved after the NOR test. In contrast, SAHA-mediated chromatin changes in TS neurons were enriched within the triplicated genomic region and were characterized by an unexpected overall decrease in chromatin accessibility (Fig. 3D, Suppl Fig. 3A-B and D).

At the transcriptional level, we detected widespread gene expression changes, with a pronounced enrichment of upregulated genes located on the triplicated chromosome (Fig. 3E, Suppl Fig. 3C, Table S5). This transcriptional imbalance closely resembled the pattern observed in trisomic animals analyzed in the home-cage condition (Fig. 1E), indicating that the dosage-driven transcriptional signature persists after the learning task. Conversely, SAHA treatment rescued transcriptional changes in TS neurons and, in line with the chromatin remodeling observed, led to a selective downregulation of triplicated genes (Fig. 3E, Suppl Fig. 3C, Table S5).

Strikingly, no DEGs were identified between WT and TS-SAHA neurons (Fig. 3E, right, Suppl Fig. 3E), indicating a complete transcriptional rescue following HDAC inhibition. This transcriptional restoration was further corroborated by analyses at the level of individual peaks and genes showing that a significant fraction of trisomy-associated DARs and DEGs were normalized by the treatment (Fig. 3 F-G). While this effect was not complete at the chromatin accessibility level, the transcriptional rescue was remarkably robust, with the vast majority of both upregulated and downregulated DEGs in TS neurons returning to baseline expression levels upon treatment.

To gain mechanistic insight into these rescuing effects, we performed a motif enrichment analysis of differentially accessible regions. Regions displaying reduced chromatin accessibility in TS neurons relative to WT were significantly enriched for binding motifs of the transcription factors FOS and JUNB (Fig 3H, Suppl Fig. 3F), which are key regulators of activity-regulated gene induction during NOR learning^35–37^. Importantly, this depletion, likely contributing to long-term memory impairments in TS mice, was reversed in SAHA-treated TS neurons, suggesting that HDAC inhibition restores the capacity of trisomic neurons to engage learning-induced transcriptional programs required for long-term memory formation.

In contrast, regions exhibiting increased accessibility in TS neurons were enriched for motifs associated with neurodevelopmental transcription factors, such as PTF1a, HEB, NEUROD1 and ASCL1 (Fig 3H, Suppl Fig. 3F), reflecting a persistent neurodevelopmental chromatin signature in trisomic neurons. Notably, this signature was also normalized following SAHA treatment (Fig. 3H).

Finally, this transcriptional rescue was further confirmed at the functional level by gene set enrichment analysis (GSEA), which revealed that most gene signatures altered in TS neurons, including pathways related to synaptic function, long-term potentiation (LTP), and learning, were reversed in SAHA-treated TS neurons (Fig 3I, Table S6), providing pathway-level confirmation that HDAC inhibition restores neuronal programs essential for cognitive function.

Altogether, our results indicate that the cognitive rescue mediated by SAHA is accompanied by substantial changes at the molecular level, including the complete normalization of the neuronal transcriptome and selective chromatin accessibility changes, consistent with a potential heterochromatinization of the triplicated genomic region.

### HDAC inhibition molecular rescue is mediated by a decreased accessibility of the triplicated region

Intrigued by the observation that SAHA-induced chromatin accessibility changes were enriched within the triplicated region, we next focused our analyses on this genomic region. As expected, a substantial fraction of genes located on the triplicated chromosome were significantly upregulated in TS neurons (Suppl Fig. 3C). Remarkably, SAHA treatment led to a robust transcriptional rescue of these triplicated genes (Fig 4A, Suppl. Fig. 4A), which was paralleled by a decreased chromatin accessibility at their associated peaks (Fig 4B, Suppl. Fig. 4A).

**Figure 4.**
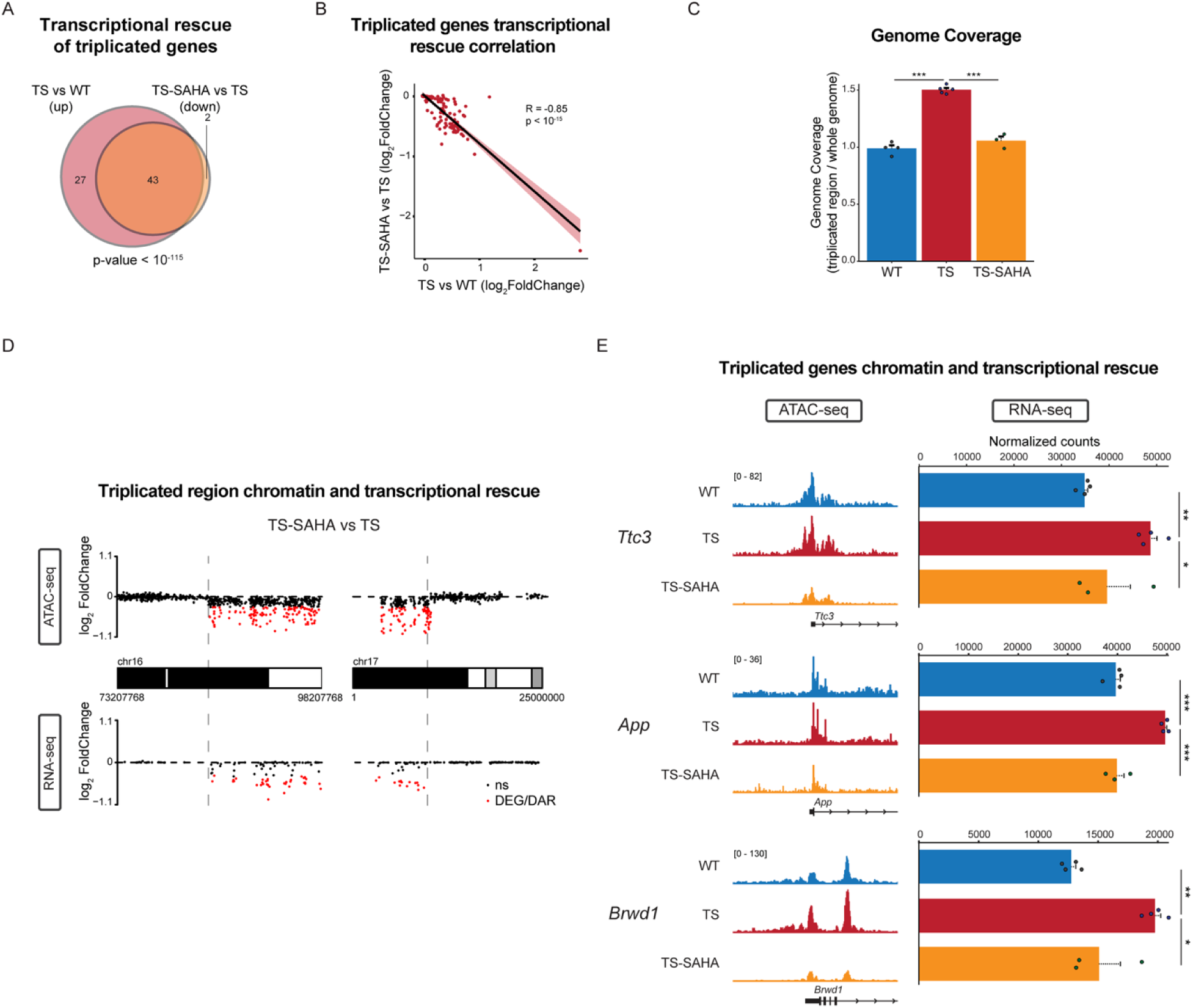
SAHA transcriptional rescue is mediated by heterochromatization of the triplicated region. **A.** Venn Diagram showing the fraction of triplicated genes whose expression is rescued by SAHA treatment. **B.** Correlation between the overexpression of triplicated genes and of normalization after SAHA treatment. **C.** Genome coverage across the triplicateda region in WT, TS and SAHA-TS neurons. **D.** Karyoplot showing the genomic loci of DARs (top, red dots) and DEGs (bottom, red dots). The dotted grey lines indicate the chromosomal breakpoints of the triplicated chromosome in Ts65Dn mice. **e.** Representative examples of heterochromatization (ATAC signal IGV tracks, left) and transcriptional rescue (right).

These results raised the intriguing possibility that SAHA induces a coordinated heterochromatization of the triplicated chromosome. To test this hypothesis, we quantified the ATAC-seq genome coverage of the triplicated region in each experimental group (see Methods). Because the Ts65Dn model carries an additional copy of this genomic region, ATAC-seq reads mapping to the triplicated interval were normalized to account for copy number differences, allowing direct comparison of chromatin accessibility across genotypes. Consistent with the presence of a supernumerary chromosome, untreated TS neurons exhibited an approximately 1.5-fold increase in genome coverage over the triplicated region compared to WT neurons (Fig. 4C). Strikingly, this increase was no longer observed in SAHA-treated TS neurons, which showed a genome coverage across the triplicated region comparable to that of WT controls. These results support the notion that SAHA promotes a global heterochromatization of the supernumerary chromosome.

Such heterochromatinization provides a mechanistic explanation for the preferential enrichment of regions with decreased chromatin accessibility within the triplicated domain and for the associated downregulation of triplicated genes following SAHA treatment (Fig. 4D; Suppl. Fig. 4A and B). This coordinated epigenetic and transcriptional rescue is exemplified at the promoters of key trisomy-associated genes, including *Ttc3*, *App*, and *Brwd1*, which show reduced chromatin accessibility concomitant with normalization of gene expression levels in SAHA-treated TS neurons (Fig. 4E).

## Discussion

DS is a disorder of gene expression deregulation, as the triplication of HSA21 results in a global disturbance of the transcriptome that is proposed to contribute to its phenotypic features. However, the mechanisms leading to this global perturbation are still not well-understood.

Here, we report that trisomic neurons show not only genome-wide transcriptional alterations, but also an altered chromatin accessibility landscape compared to their euploid counterparts, characterized by a loss of accessibility at gene promoters. This result is in line with our previous findings in trisomic hippocampi, where we detected a global decreased histone acetylation at H3K9 and H3K14 residues^11^, as these modifications are generally associated with active promoters^38–40^. Notably, H3K14 acetylation is known to mark inactive but inducible, or primed, promoters^39^. Therefore, its reduction in trisomic neurons could impair the primed state of gene promoters, reduce chromatin accessibility, and ultimately hinder gene expression during learning. Among the promoters showing reduced accessibility in our work, are genes critical for neuronal plasticity and hippocampus-dependent learning, including *Homer1*, *Snx27*, and *Numb*^29–32^. These findings are consistent with previous work linking cognitive dysfunction to reduced histone acetylation around TSSs^24^.

Given the reversible nature of histone acetylation, we sought to rescue its deficits using the clinically-approved HDACi SAHA. Chronic SAHA treatment has previously been shown to improve molecular phenotypes of a Huntington’s disease mouse model^41^ and to restore spatial memory in mouse models of age-associated memory impairment^42^. Here, we report that SAHA treatment fully rescues cortico-hippocampal memory deficits in TS mice as assessed by NOR, a paradigm in which TS mice are particularly impaired^33,34^.

Unexpectedly, our data reveal that histone acetylation deficits in TS mice are only partially corrected by SAHA, suggesting that the mechanisms underlying behavioral rescue are more complex than initially hypothesized. Integrated analysis of transcriptional and chromatin accessibility profiles in treated and untreated mice following the NOR learning task revealed the surprising nature of this rescue. Strikingly, SAHA treatment leads to a near-complete normalization of the transcriptional alterations in TS neurons. At the chromatin accessibility level, however, its genome-wide effects were more limited and appeared largely restricted to the triplicated chromosomal region. In particular, SAHA treatment was associated with reduced chromatin accessibility within the genomic region that is triplicated in the Ts65Dn mouse model. This normalization of the overexpression of the triplicated genes and this likely underlies the broader genome-wide transcriptional alterations.

At first glance, this observation appears counterintuitive, as HDAC inhibition is typically expected to increase histone acetylation and thereby promote chromatin accessibility and gene expression. However, it aligns with previous studies suggesting that HDACis can modulate transcription, selectively normalizing dysregulated genes, rather than inducing widespread expression changes^43,44^. Although the mechanisms driving the specificity of HDACi effects in the brain remain unclear, a study in breast cancer reported that HDACis preferentially repress the transcription of high copy number genes by blocking RNA polymerase II elongation^45^. Additionally, a recent study indicates that HDACi-induced gene repression can occur without the gain of repressive chromatin marks and may instead involve repressive chromatin looping^46^. Similar mechanisms could potentially contribute to the normalization of extra gene copies in trisomic neurons observed here, although this possibility will require further investigation. Finally, we did not detect clear enrichment of specific transcription factor motifs within the triplicated region showing decreased accessibility following SAHA treatment (data not shown), suggesting that the observed chromatin compaction is unlikely to be driven by the recruitment of a single dominant transcriptional repressor. Instead, these findings are more consistent with a broader structural reorganization of chromatin within the supernumerary chromosomal region.

## Limitations of the study

Several limitations of this study should be considered when interpreting our findings. First, although our results suggest that SAHA treatment is associated with reduced chromatin accessibility within the triplicated genomic region, the molecular mechanisms leading to this apparent heterochromatinization remain unclear and require further investigation.

Another limitation relates to the interpretation of chromatin accessibility changes following the behavioral task. In this case, differences in chromatin accessibility may partially reflect variation in neuronal activation states induced by the NOR task rather than purely genotype- or treatment-driven chromatin remodeling. Future studies combining chromatin accessibility profiling with single-cell or activity-tagging approaches may help to further disentangle activity-dependent chromatin changes from constitutive epigenetic alterations associated with trisomy.

Finally, among the Down syndrome (DS) mouse models generated so far, only Ts65Dn, Tc1, Ts66Yah, and TcMAC21 represent true aneuploid models carrying a freely segregating supernumerary chromosome. This feature may be relevant for modeling dosage-driven chromosomal effects. For our study, we chose the Ts65Dn model because it reproduces many key characteristics of DS. However, it is important to note that Ts65Dn mice also harbor a triplication of 43 protein-coding genes that are not orthologous to HSA21 and are not triplicated in individuals with DS. This additional genetic content may influence some of the molecular phenotypes observed in this study.

## Resource availability

### Lead contact

Requests for additional information, resources, and reagents should be directed to and will be fulfilled by the corresponding authors, Cesar Sierra and Mara Dierssen.

### Materials availability

This study did not generate new reagents.

### Data and code availability

- The ATAC-seq RNA-seq data that support the findings of this study have been deposited in European Nucleotide Archive (ENA), with the accession number PRJEB110155. All additional raw and processed data and supplementary materials are available upon request.
- The code used in this manuscript to analyze the ATAC-seq and RNA-seq data can be found in: https://github.com/cesarsierran/TS_SAHA_omics

## Acknowledgments

The lab of MD is recognized by the Secretaria d’Universitats i Recerca del Departament d’Economia I Coneixement de la Generalitat de Catalunya (Grups consolidats 2023). We acknowledge the support of H2020 SC1 Gene overdosage and comorbidities during the early lifetime in Down Syndrome GO-DS21-848077, Agencia Estatal de Investigación (PID2022-141900OB-I00 INTO-DS), Fundació La Marató-TV3 (#202212-30), Ministerio de Ciencia Innovación y Universidades (RTC2019-007230-1, RTC2019-007329-1; and CPP2022-009659).. We acknowledge the support of the Spanish Ministry of Science and Innovation through the Centro de Excelencia Severo Ochoa (CEX2020-001049-S, MCIN/AEI/10.13039/501100011033), the Generalitat de Catalunya through the CERCA program and EMBL partnership. The CIBER of Rare Diseases is an initiative of the Instituto Carlos III (ISCIII). CS received the FI grant from Agència de Gestió d’Ajuts Universitaris i de Recerca (AGAUR) de la Generalitat de Catalunya. RAJC was supported by the Jérôme-Lejeune Foundation with the Sisley d’Ornano-Lejeune postdoctoral fellowship 2021 (Grant Number: 15_PDC-2021) and the Sello Excelencia ISCIII-Health (Grant Number: IHMC22/00026).

## Author contributions

**MD** and **CS** conceived, designed and coordinated the study, and wrote the manuscript. **CS** collected and analyzed sequencing datasets. **CS** conducted the behavioural experiments. **RAJC** performed the Western blot quantification. All authors revised and corrected the final version of the manuscript. Correspondence should be addressed to **MD** and **CS**.

## STAR⍰Methods

### Animals

4-months old Ts(17(16))65Dn (Ts65Dn; TS), which contains two thirds of genes orthologous to HSA21, and their wild type littermates were used. Experimental mice were generated by crossing of TS females to C57/6Ei × C3H/HeSnJ F1 hybrid (B6EiC3) males. The parental generation was obtained from the research colony at the Jackson Laboratory (B6EiC3Sn.BLiA-Ts(1716)65Dn/DnJ; Stock No: 005252). Genotypes of mice were authenticated by PCR assays on mouse tail samples with an in-house protocol. Mice were housed in standard cages (156 x 369 x 132 mm), with food and water available ad libitum in standard conditions (12:12 light cycle; 400 lux). Sawdust and nesting materials in each cage were changed once a week, but never on the day before or the day of testing, to minimize the disruptive and stressful effect of cage cleaning on behavior.

All experiments followed the principle of the “Three Rs”: replacement, reduction and refinement according to Directive 63 / 2010 and its implementation in Member States. The study was conducted according to the guidelines of the local (law 32/2007) and European regulations (2010/63/EU) and the Standards for Use of Laboratory Animals no. A5388-01 (NIH) and approved by the Ethics Committee of Parc de Recerca Biomèdica (Comité Ético de Experimentación Animal del PRBB (CEEA-PRBB); MDS 0035P2). The CRG is authorized to work with genetically modified organisms (A/ES/05/I-13 and A/ES/05/14). Experimenters were blinded to the genotype for all experiments. Sample size was determined based on previous ATAC-seq and RNA-seq experiments and behavioural studies that demonstrated sufficient statistical power.

### SAHA administration

The HDACi SAHA was dissolved in drinking water following a previously published formula^41^: 0.67g of SAHA was dissolved in 1L of drinking water containing 18g of HP-β-CD (Sigma-Aldrich #332607). The solution was heated to 95°C and stirred until no SAHA particles were visible (approximately 20 min) and then allowed to cool down to room temperature before making it available to animals. In an initial test, water intake of individualized animals was monitored to assess whether the potential taste of the drug was affecting it. Animals’ liquid intake was on average 3 mL/day regardless of the presence of the drug, which equals 2 mg SAHA per day. Treatment with SAHA started at P90 and was finalized at P120 (Fig. 2). Weight was monitored along the treatment to discard any toxicity derived from SAHA administration. Drinking water containing 18g of HP-β-CD/L was used as control.

### ATAC-seq and RNA-seq experiments

#### Fluorescence-Activated Nuclear Sorting (FANS)

The hippocampus of each mouse was dissected and placed in chilled Hanks’ Balanced Salt Solution (HBSS; #H6648). To obtain a nuclei suspension, each hippocampus was transferred to a new tube containing 500 μL chilled EZ Lysis Buffer (#NUC-101) and homogenized using a sterile RNase-free douncer. The resulting homogenate was filtered using a 70 μm-strainer mesh to remove remaining chunks of tissue and centrifuged at 500 g for 5 min at 4°C. The pellet containing the nuclei was resuspended in 1.5mL EZ Lysis Buffer and centrifuged again. The supernatant was removed and 500 μL of Nuclei Wash and Resuspension Buffer (NWRB, 1X PBS, 1% BSA and 0.2 U/μL RNase inhibitor #N8080119) were added without disturbing the pellet and incubated for 5 min. After incubation, 1 mL of NWRB was added and the pellet resuspended. The nuclei suspension was centrifuged again and the washing step was repeated with 1.5 mL of NWRB. After an additional centrifugation, nuclei were resuspended in 500 μL of 1:1000 anti-NeuN antibody conjugated with AlexaFluor 647 (Abcam #ab150115) in PBS and incubated in rotation for 15 min at 4°C. After incubation, nuclei were washed with 500 μL of NWRB and centrifuged again. Last, nuclei were resuspended in NWRB supplemented with DAPI and filtered with a 35 μm cell strainer to obtain a single-nuclei suspension. This suspension was then used to sort by FANS the neuronal nuclei that were used for ATAC-sequencing and RNA-sequencing.

#### ATAC-seq experiment

To perform ATAC-seq, 50,000 FANS-sorted nuclei were pelleted at 4 °C, 1,000 × g for 7 min, resuspended in 50 μl of transposition mix (25μL buffer + 22.5μL H2O + 2.5μL Tn5) and gently pipetted to resuspend the nuclei. Transposition reaction was incubated at 37°C for 45 min in a thermomixer with shaking at 1000 r.p.m. The transposed DNA was purified using GenElute PCR Clean-up (Sigma; #NA1020). To reduce PCR-induced biases during library preparation, after the first 5 PCR cycles, 10% of the sample was monitored with SYBR-green qPCR for 20 cycles to calculate the 25% of saturation per sample (every sample needed more than 7 PCR cycles in total). Final libraries and purification were performed by two-side size selection using magnetic beads.

Sequencing was performed using an Illumina HiSeq2500 apparatus at a depth of at least 80 million reads per sample.

#### ATAC-seq data processing and downstream analyses

After trimming adaptors, 50 bp paired-end reads were aligned using bowtie2 (v.2.2.6) to the mouse reference genome (GRCm38/mm10). Duplicated reads were removed using Picardtools (v.2.17) and only paired reads with very high mapping quality (MAPQ score > 30) were used for further analysis. In order to account for the triplicated region in Ts65Dn, reads mapping to this region were divided by 1.5 in trisomic samples. ATAC-seq peak regions of each sample were called using MACS2 with parameters -q 0.05 --nomodel --extsize 200 --broad. Regions listed in the ENCODE Blacklist were excluded from called peaks.

Following this preprocessing step, ATACseqQC package^47^ was employed to perform Quality Control of each individual sample. Low quality samples (fraction of reads in peaks, FRiP < 0.1) were discarded for downstream analyses.

A MA plot was then used to visualize changes in chromatin accessibility for all peaks. For ATAC-seq peaks common to at least two samples, read counts were normalized by dividing the raw reads by the FRiP value. We then accessed the significant change of chromatin accessibility between different groups using DESeq2. It was defined as significantly changed if the peak showed FDR < 0.1.

#### Transcription factor motif enrichment

For each comparison, significant peaks were tested for overrepresentation of each DNA motif using HOMER^48^. The resulting p values were FDR adjusted.

#### RNA-seq experiment

For RNA-seq, experiments were performed in parallel to ATAC-seq at P120. 200,000 nuclei were FANS-sorted sorted directly in RLT Buffer for purification using the RNeasy Micro Kit (Qiagen #74004). RNA integrity (RIN) was checked by 2100 Bioanalyzer and only samples with a RIN > 8 were further processed. Library preparation and sequencing were performed by the Genomics facility of the Center for Genomic Regulation (CRG, Barcelona). Libraries were prepared using the SMARTer Stranded Total RNA-Seq Kit v2 from ribo-depleted total RNA. Samples were sequenced (paired end, 50 base pairs in length) using an Illumina HiSeq2500 apparatus with a depth of at least 50 million reads.

Reads were then aligned using STAR (v.2.5.2a)^49^ to reference genome (mm10) and quantified using Rsubreads (v.1.26)^50^. Gene expression data was visualized by PCA plot and one WT sample in the naïve experiment was identified as an outlier and removed for downstream analyses.

Differential expression analysis was performed using DESeq2 (v.1.10.0)^51^. Genes with FDR < 0.05 were considered significantly differentially expressed.

#### Gene set enrichment

Differentially expressed genes in each comparison were tested for enriched Gene Ontology processes, using a hypergeometric test (shinyGO^52^), and corrected for multiple hypotheses by FDR. Processes with p adjusted value < 0.05 were reported as significantly enriched. The complete list of genes detected in the dataset was used as the universe for the hypergeometric test.

#### Karyoplots of RNA and ATAC-seq data

The karyoplots indicating the localization of DARs and DEGs in the whole genome were created using the R package karyoploteR^53^.

### Western Blotting for histone acetylation

To determine histone acetylation, hippocampal tissues were dissected and histones isolated from the whole hippocampus by acid-extraction. Hippocampi were placed in homogenization buffer (50 mmol/L Tris-HCl, pH 7.5, 25 mmol/L KCl, 250 mmol/L sucrose, 2 mmol/L sodium butyrate, 1 mmol/L sodium orthovanadate, 0.5 mmol/L PMSF, 1X protease inhibitor cocktail). The homogenate was then centrifuged at 7700g for 1 min at 4°C. The pellet containing the nuclear fraction was resuspended in 500 μL of 0.4N H2SO4 by thorough trituration, then incubated on ice for 30 min with intermittent vortexing. Then, these 1.5 mL tubes were centrifuged at maximum speed for 10 min at 4°C. The supernatant was transferred to new tubes, and the histones were precipitated with 250 μL of trichloroacetic acid and sodium deoxycholate on ice for 30 min, then centrifuged at maximum speed for 30 min at 4°C. The pellet was rinsed with acetone, centrifuged again, and then dried for 5 min. Nuclear histone protein samples were resuspended in 50 μL of 10 mM Tris-HCl (pH 8.0). Protein concentration was measured by Pierce BCA Protein Assay Kit (Thermo Fisher #23225). A total 1 μg was loaded with Laemmli buffer in each lane of a 15% acrylamide gel. The histones of each hippocampus were loaded in two lanes as duplicates. The fractionated histones were transferred to PVDF membranes using the semi dry iBlot 2 (Thermofisher #IB21001) system. The membranes were then blocked with 5% milk in TBS-Tween 0.1% (TBS-T) for 1 hour before immunodetection with anti-diacetyl lysine 9 and lysine 14 histone H3 (AcH3K9K14) antibody (Cell Signaling Technology #9677) at a 1:1000 dilution and with anti-histone H4 (Millipore #07-108) at a 1:500 dilution. Primary antibody incubation was followed by three 10 min washes at RT in TBS-T before incubation with IRDye 800CW Goat anti-Rabbit IgG antibody (LI-COR Biosciences #925-32211), three washes and visualization using Odyssey Platform (LI-COR). Western blot’s bands were quantified by densitometry using ImageStudio. The total amount of H4 protein was used as the loading control. Independent western blotting experiments were integrated by median normalization, excluding extreme values and statistical differences in the ratio AcH3/H4 were assessed with a linear mixed-effect model allowing for nested random effects (for samples coming from the same mouse).

### Novel Object Recognition

The NOR protocol consists of three phases, namely habituation, familiarization and recall. During the habituation session, mice were allowed to freely explore the empty open field for 10 minutes. 24 hours later, each mouse was returned to the arena containing two identical objects placed at symmetrical positions 5 cm from the arena wall and allowed to explore them freely for 15 minutes. During the familiarization session most mice reached a minimum exploration for each object of 30 seconds. Mice not reaching this threshold, were excluded from the analysis. After a retention interval of 24 hours, the mouse was returned to the arena in which one of the objects was replaced by a novel object and let to explore both the familiar and the novel object for 5 minutes. All sessions were video-taped.

#### NOR behavioural experimental variables

Locomotor activity was quantified as the total distance traveled in the apparatus during the experimental sessions. Thigmotaxis refers to the disposition to remain close to the walls of the apparatus. It is measured as distance traveled or percentage of time spent in the periphery of the apparatus. It decreases gradually during the first minutes of exploration, and can be used as an index of anxiety^54^. Exploration time is defined as the action of pointing the nose toward an object, at a maximum distance of 2 cm or touching it. Going around the objects or sitting on the object is not evaluated as exploration time. As such, the exploration time is only computed when the snout of the animal is directed toward the object, sniffing or touching it. Discrimination index (DI), which is a measure of the recognition memory of the animal, is calculated as the difference in exploration time for the familiar versus the novel object, divided by the total amount of exploration.

#### Statistical analyses of behavioral and WB experiments

To investigate statistical differences in experimental variables, two-way ANOVA was conducted. For pairwise comparisons, the Saphiro-Wilk and Bartlett tests were used to assess normality of values and homogeneity of variances between groups, respectively. When the two conditions were met, pairwise differences were tested by Tukey’s post hoc test. All statistical analyses were two-tailed. P values were considered to be significant when α < 0.05.

## Supplementary Figures

**Supplementary Figure 1.**
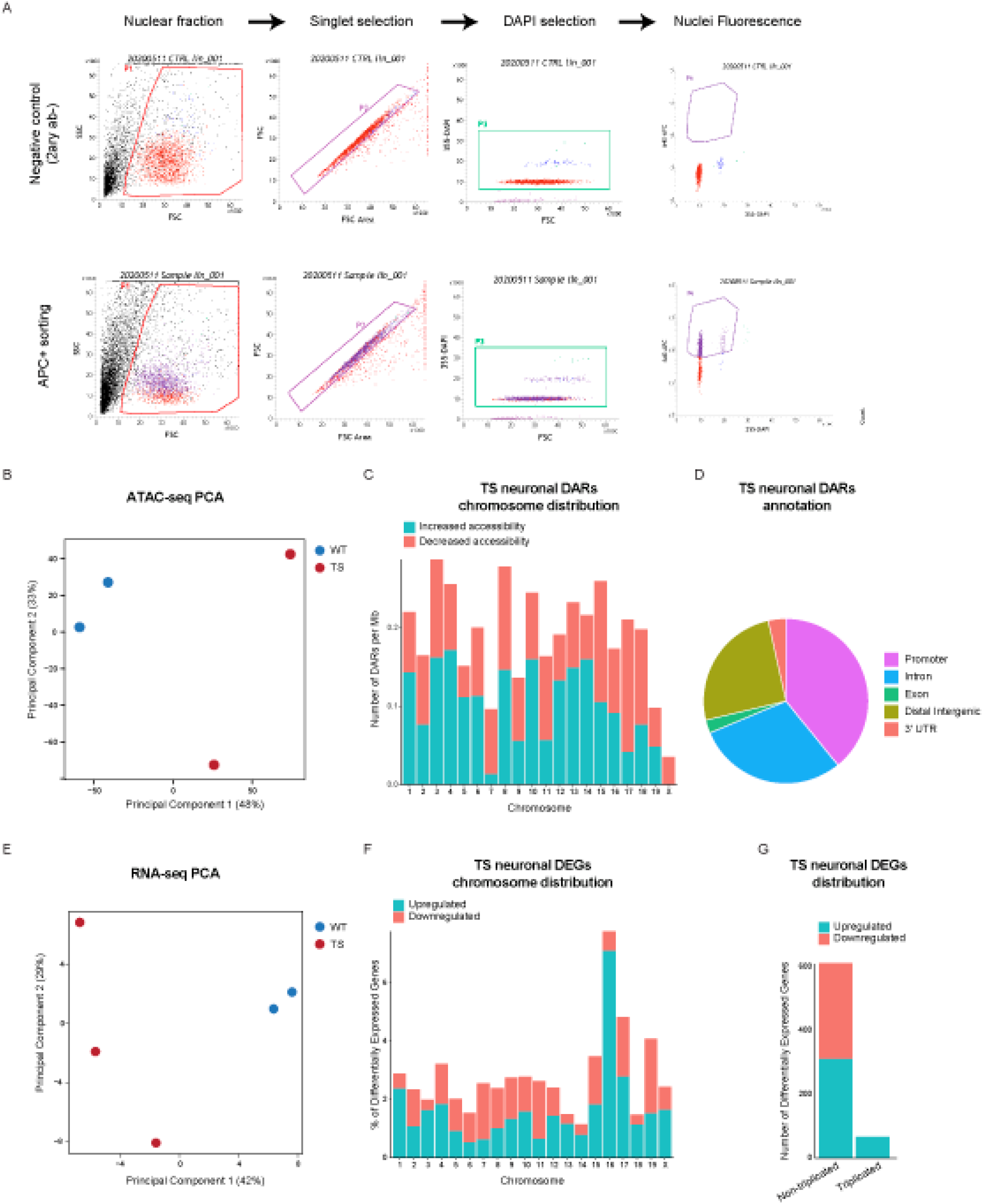
Trisomy induces broad changes in the transcriptome and epigenome. **A**. Step by step analysis and channel filtering (selected filtered population indicated) of flow cytometry signal for the isolation of fluorescent singlet nuclei. Comparison of a control (no antibody) sample (above) vs anti-NeuN antibody sample (below) mice. Similar results were obtained for each biological replicate. **B.** PCA plot of ATAC-seq data obtained from two WT and two TS mice. See Methods. **C.** Proportion of differentially accessible regions in TS neurons located in each chromosome (number of DARs / Mb) **D.** Proportions of DARs annotated to different genomic regions. **E.** PCA plot of RNA-seq data obtained from two WT and three TS mice. See Methods. **F.** Proportion of genes located in each chromosome that were found differentially expressed in TS neurons. **G.** Barplot showing the number of differentially expressed genes by their triplication status.

**Supplementary Figure 2.**
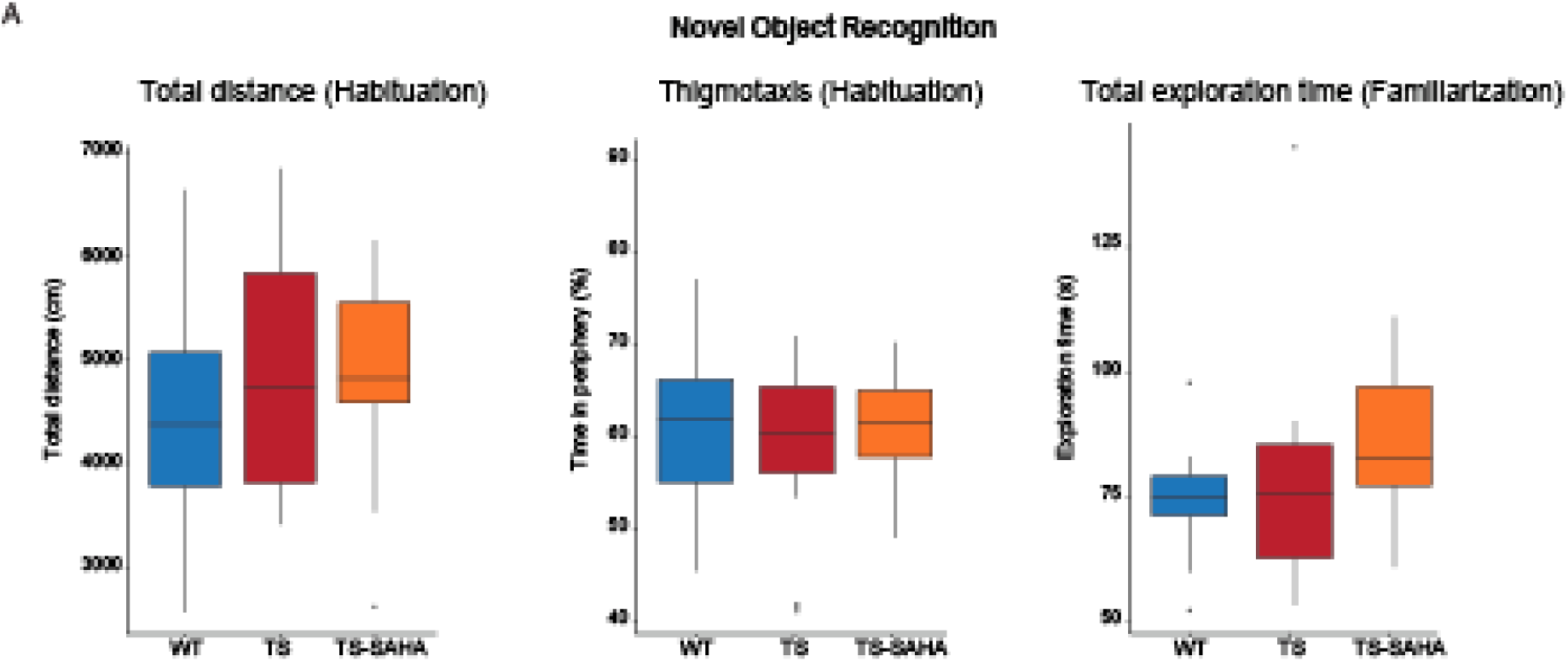
Novel Object Recognition experimental variables are unaltered by SAHA treatment. **A.** Box plots depicting total distance travelled (left), thigmotaxis (center) and exploration time (right) in the four experimental groups. WT n = 40, TS n = 30, TS + SAHA n = 30. Two-way ANOVA, Tukey HSD as Post-hoc.

**Supplementary Figure 3.**
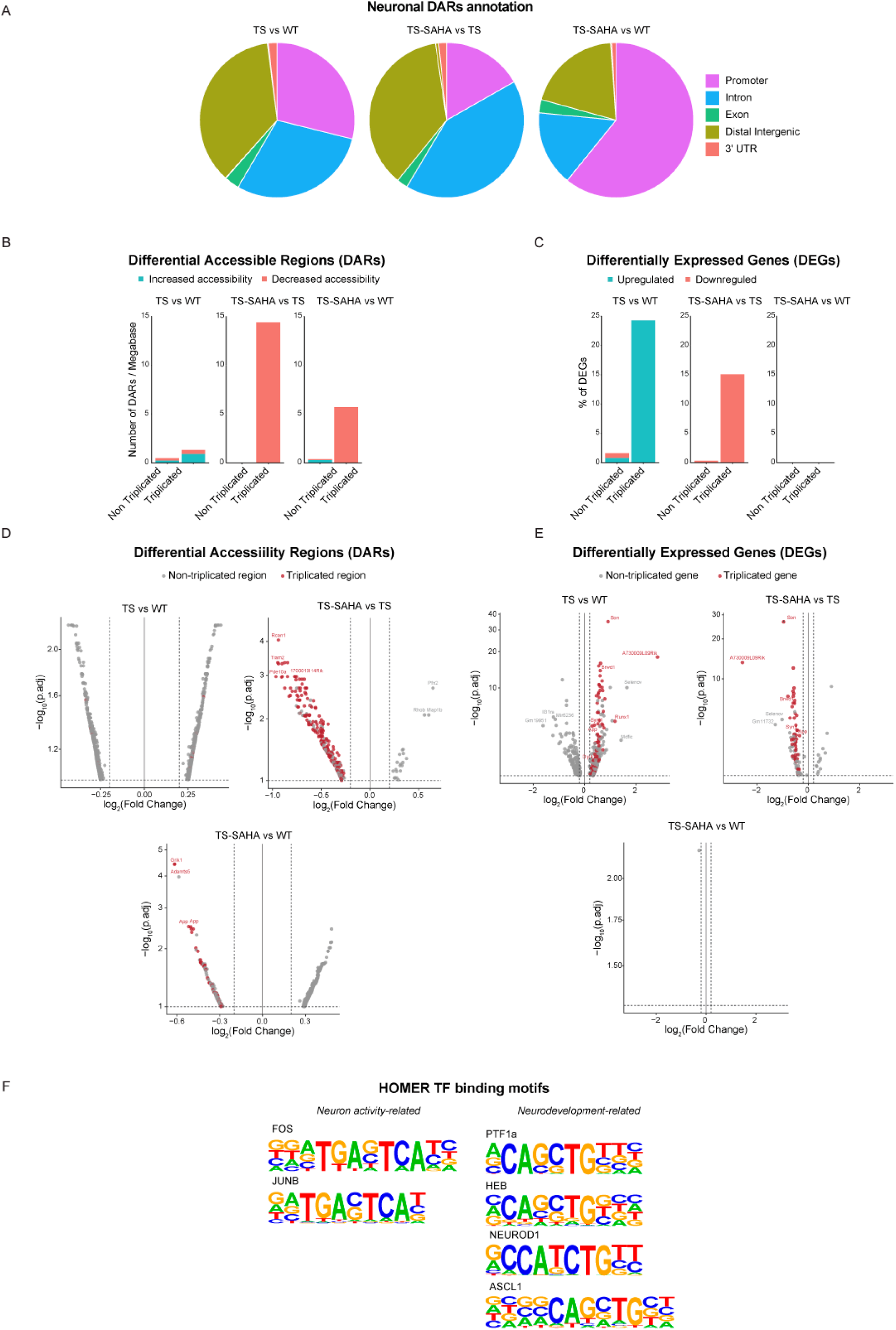
Molecular characterization of SAHA-mediated changes. **A**. Proportions of DARs annotated to different genomic regions (promoter, intron, exon, distal intergenic and 3′ UTR) for each comparison (TS vs WT, TS-SAHA vs TS, and TS-SAHA vs WT). **B.** Barplots showing the number of DARs per megabase in the non-triplicated and triplicated genomic regions. **C.** Barplots showing the proportion (%) of differentially expressed genes according to their triplication status. **D.** Volcanoplots showing the DARs for each comparison **E.** Volcanoplots showing the DEGs for each comparison. **F.** Binding motifs of transcription factors enriched among DARs, including neuronal activity-related (FOS, JUNB) and neurodevelopment-related (PTF1a, HEB, NEUROD1, ASCL1) factors.

**Supplementary Figure 4.**
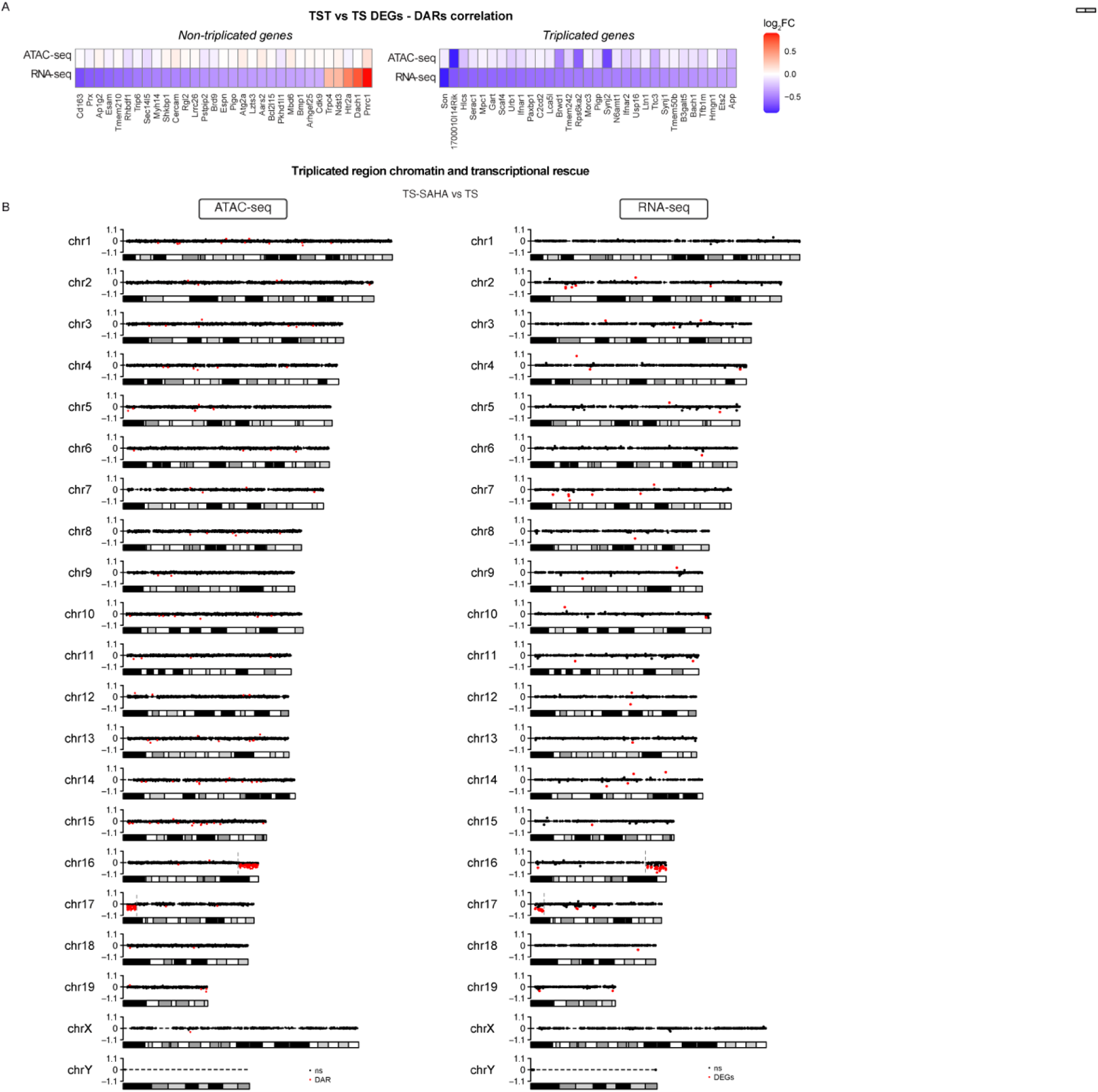
Trisomy-specific impact of SAHA treatment. **A**. Heatmap showing transcriptional and chromatin accessibility changes in non-triplicated (left) and triplicated (right) DEGs. **B.** Karyoplot showing the genomic loci of DARs (left, red dots) and DEGs (right, red dots)

